# Continuity of Mitochondrial Budding: Insights from BS-C-1 Cells by *in-situ* Cryo-Electron Tomography

**DOI:** 10.1101/2023.11.10.566563

**Authors:** Judy Z Hu, Lijun Qiao, Xianhai Zhao, Chang-Jun Liu, Guo-Bin Hu

## Abstract

Mitochondrial division is a fundamental biological process that is crucial to cellular functionality and vitality. The prevailing hypothesis of Drp1 regulation with the involvement of ER and cytoskeleton does not account for all the observations. Following up our previous study in HeLa cells which led to the new hypothesis of mitochondrial division by budding, we employed *in-situ* Cryo-Electron Tomography (Cryo-ET) to visualize mitochondrial budding in intact healthy monkey kidney cells (BS-C-1 cells). Our findings reaffirm single and multiple mitochondrial budding, supporting the new hypothesis. Notably, the budding regions vary significantly in diameter and length, which may represent different stages of budding. More interestingly, no rings, or ring-like structures, or ER wrapping is presented in the budding regions suggesting mitochondrial budding is independent from Drp1 and ER. Meanwhile, we uncovered direct interactions between mitochondria and large vesicles, distinct from small mitochondrial-derived vesicles and extracellular mitovesicles. We propose these interacting vesicles may have mitochondrial origins.

## Introduction

Mitochondria are considered the power plants for eukaryotic cells (Voet et al., 2016). They are also related to many diseases such as cancer (Douglas 2012), Alzheimer’s disease (Moreira 2010), and Parkinson’s disease (Exner et al. 2012). New mitochondria are produced only by division of existing mitochondria; thus the success of mitochondrial division can determine the life or death of eukaryotic cells. Based on previous studies, scientists believe that mitochondria divide by Drp1- and endoplasmic reticulum (ER)-mediated fission with the involvement of cytoskeleton, which includes microtubules, actin filaments, and septin filaments (Chan 2006, Friedman et al. 2011, Mears et al. 2022, Mageswaran et al. 2023). However, the prevailing model does not account for some important observations. For example, mitochondrial fission may occur very frequently (Beech et al., 2000), which one presumes should lead to a large increase in the number of mitochondria and/or the total mitochondrial volume and mass with the increase of fission cycles as the cell division does. But no such an increase of mitochondrial volume or mass was observed after many cycles of mitochondrial fission (Pesce & Walter, 2007), which indicates mitochondrial fission may not produce new mitochondria instead it may just split mitochondria into smaller pieces. Meanwhile, it was reported that Drp1 promotes mitochondrial fusion instead of fission (Zhao et al., 2011). And mitochondrial fission was observed to occur independently of Drp1 (Helle et al. 2017). The inconsistency may be due to the technical limitations in previous studies.

Previous studies have relied heavily on visualization by conventional imaging techniques, such as fluorescence microscopy-based imaging techniques and conventional electron microscopy. However, even the latest fluorescence microscopy-based imaging technique is limited in its spatial resolution to ∼100 nm (Guo et al., 2018), which is insufficient to distinguish mitochondrial division, which involves contacts of 10-30 nm spacing (Wu et al., 2018). Moreover, fluorescence microscopy relies on labeling, which may cause researchers to miss critical structures that are not labeled. Conventional electron microscopy provides higher resolution but suffers artifacts in specimen preparation, including chemical fixation, dehydration, heavy metal staining (Kilpatrick et al., 1985), which could have resulted in ambiguous or misleading interpretation as revealed in the desmosome studies (He et al., 2003, Al-Amoudi et al., 2007). To overcome the limitations of conventional imaging techniques in the studies of mitochondrial division, one desires an artifact-free imaging technique with sufficient resolution.

The emerging *in-situ* Cryo-Electron Tomography (Cryo-ET) is a technique that provides not only resolution better than 10 nm (Turk and Baumeister 2020) but artifact-free specimen preparation without the need of labeling. Furthermore, it provides three-dimensional views. We previously employed Whole Cell Cryo-ET (now a part of *in-situ* Cryo-ET) to visualize mitochondria in frozen hydrated intact HeLa cells (Hu 2014), in which we observed many mitochondrial morphological and structural features that resemble the budding alpha-proteobacteria (Hirsch 1974). Interestingly, based on genomics, mitochondria are believed to have originated by permanent enslavement of purple non-sulphur bacteria (Cavalier-Smith 2006), which belong to alpha-proteobacteria. Therefore, we proposed a new hypothesis for mitochondrial division, which is that mitochondria divide by budding (Hu 2014).

An immediate question is whether the budding features observed in HeLa cells are HeLa specific, since HeLa cells are cancer cells, or generally applicable to other types of cells. In this study, we carried out the investigation in monkey kidney cells (BS-C-1) which are healthy animal cells. Our results of this study represent a general mechanism for mitochondrial division.

## Results and Discussion

Visualization of mitochondria and mitochondrial dynamics, such as mitochondrial division, has been a challenging task. For *in-situ* Cryo-ET imaging of cellular structures, it is important to prepare a thin enough specimen. With intact animal cells grown on electron microscopic grids, the cells need to have considerable flat and thin periphery regions. Fig.1 shows a small area of specimen on an EM grid we prepared in this study. The dark blobs (blue arrows) shown in the grid squares are cells. Animal cells are usually too thick for an electron-microscope beam to penetrate, which is why they look dark. Nevertheless, the growing edges or peripheries of the cells are flat and thin (green arrow), therefore bright enough for electron microscopy imaging. Data in this paper was acquired from thin cellular regions like the one indicated by the green arrow.

**Fig. 1.**
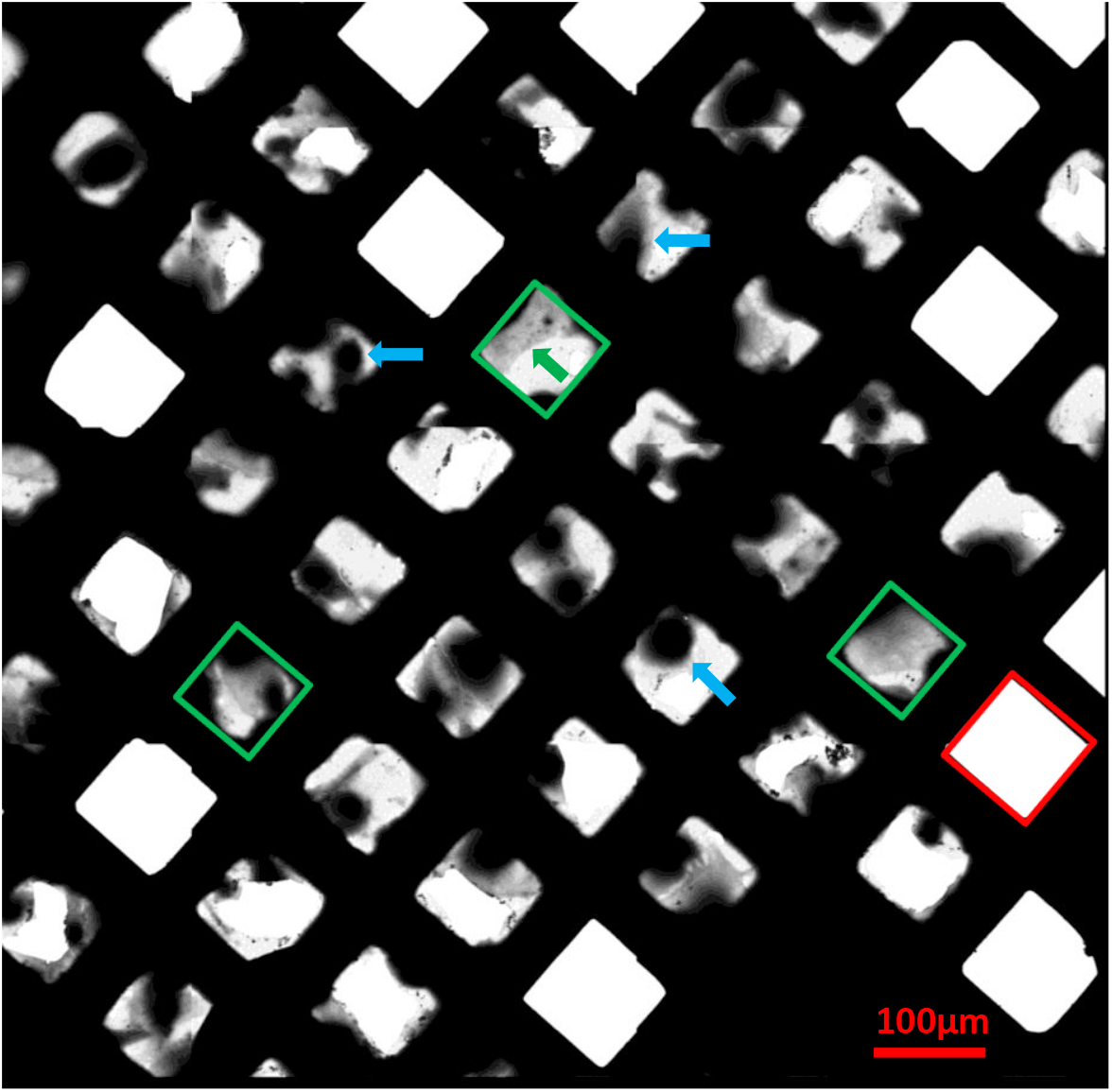
A low magnification (135x) Cryo-EM image of cells grown on electron microscopic grid and directly frozen to liquid nitrogen temperature. Three grid squares marked in green are examples of cells with peripheries (green arrow) that are suitable for high resolution *in-situ* Cryo-EM imaging. There are also empty squares (one example marked in red) where carbon support film broke and got lost during grid handling.

In *in-situ* Cryo-EM/ET, searching and locating targets for data collection is critical for data collection. A common approach for target search is to use correlative microscopy in which the targets were labeled with fluorescence signals, located by fluorescence microscopy, and then the locations are transferred to the electron microscope or tomography software for Cryo-ET data collection. In this study, we used images of intermediate magnification as shown in Fig.2 for target search and the locations only need to be aligned by the same tomography software with image shift for Cryo-ET data acquisition. The image shown in Fig.2 was taken from one of the cell peripheries shown in green boxes in Fig.1. Mitochondria and mitochondrial budding features such as narrow connections one can visualize between mitochondria (yellow arrows), make it possible to specify targets.

**Fig. 2.**
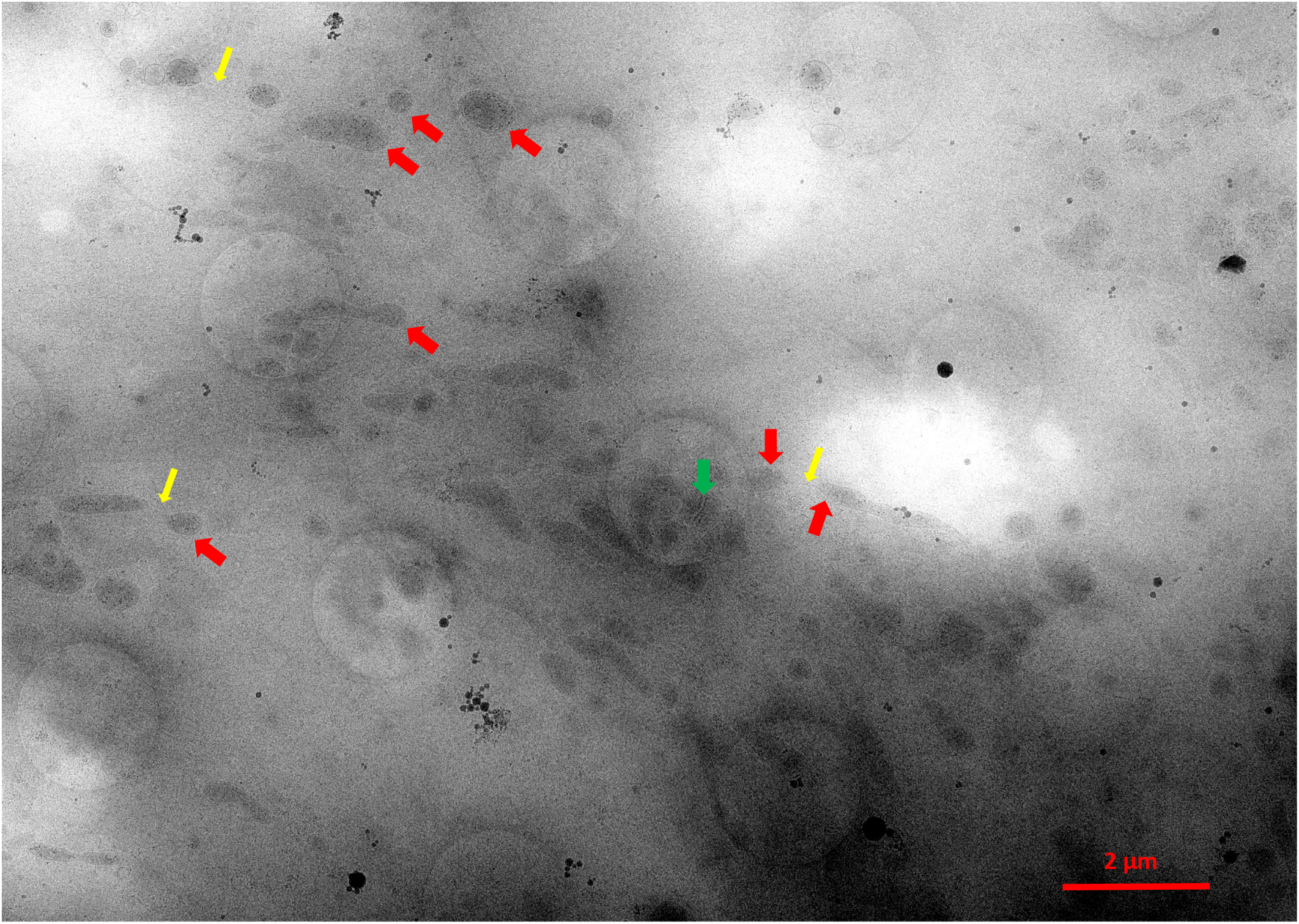
Intermediate magnification (2250x) Cryo-EM image of a cellular region for data acquisition targeting. A lot of mitochondria are presented, a few of which are indicated by red arrows. Cristae inside mitochondria can be appreciated (green arrow). Thin connecting structures (yellow arrows) are visible.

Traditionally, mitochondria were observed to have bean-like morphology, but lately scientists have observed more complicated mitochondrial morphology, such as mitochondrial networks, indicating the complexity of mitochondrial structures and functions. Compared to visualization by fluorescence microscopy which can show only the limited structures that are labeled (Guo et al., 2018), *in-situ* Cryo-ET displays everything in the cellular area imaged, as shown in Fig.3. One sees mitochondrial double membrane, mitochondrial cristae, and dark granules which are commonly observed by *in-situ* Cryo-ET (Wu G et al., 2023), also ER, microtubules (MT), and vesicles (Vs).

**Fig. 3.**
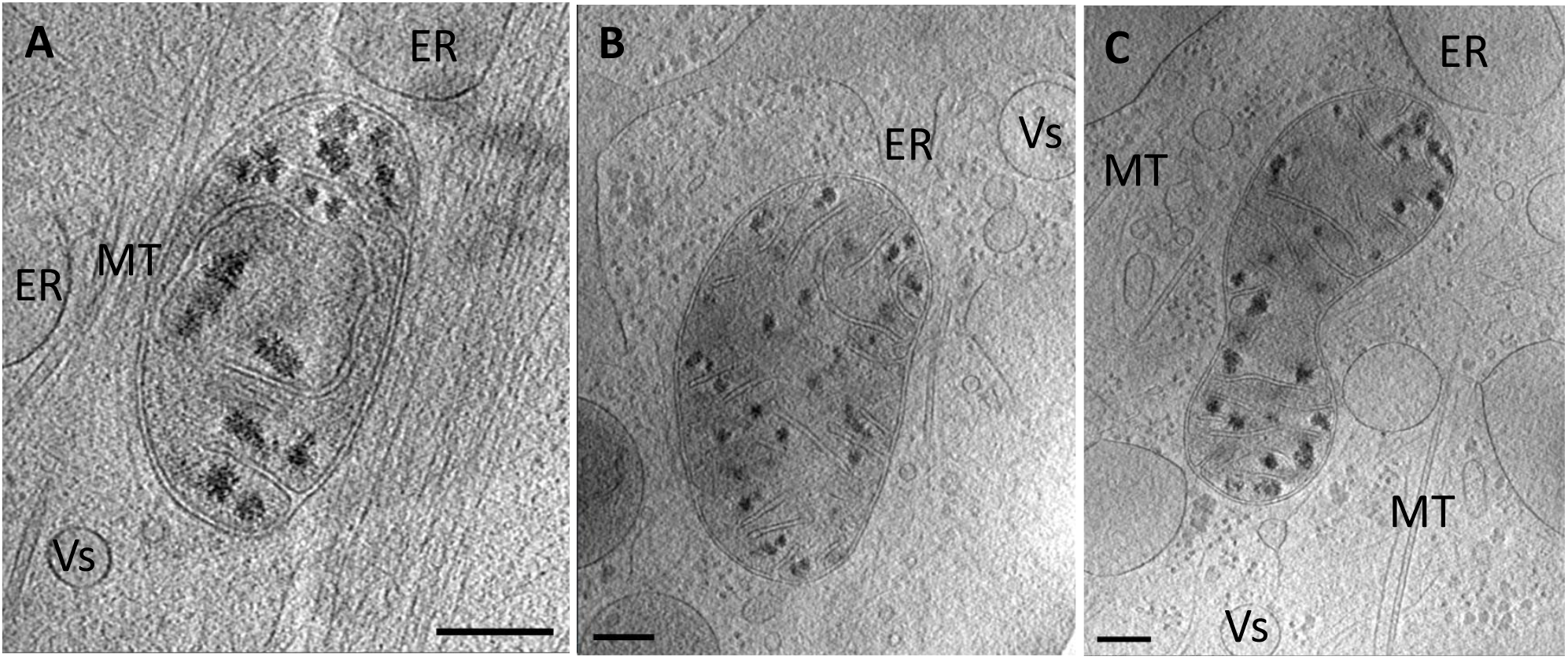
2D slices of 3D *in-situ* Cryo-electron tomography reconstruction of individual/independent mitochondria. They are accompanied by but remain some distance from endoplasmic reticulum (ER). Bars, 200nm.

Besides individual mitochondria, mitochondrial budding features were observed not only with single budding shown in Fig.4 but also multiple budding as shown in Fig.5. Many images do not show the entire mitochondria bodies because of the camera’s size limit. The budding features are very similar to what we observed in HeLa cells. And they also resemble those of the budding alphaproteobacteria (Hirsch 1974), perhaps revealing the preserved reproductive mechanism of engulfed bacteria of the endosymbiotic hypothesis. (Roger et al., 2017). Budding mitochondria may eventually form a mitochondrial network if daughter mitochondria do not split from mother mitochondria, and the budding process keeps going on.

**Fig. 4.**
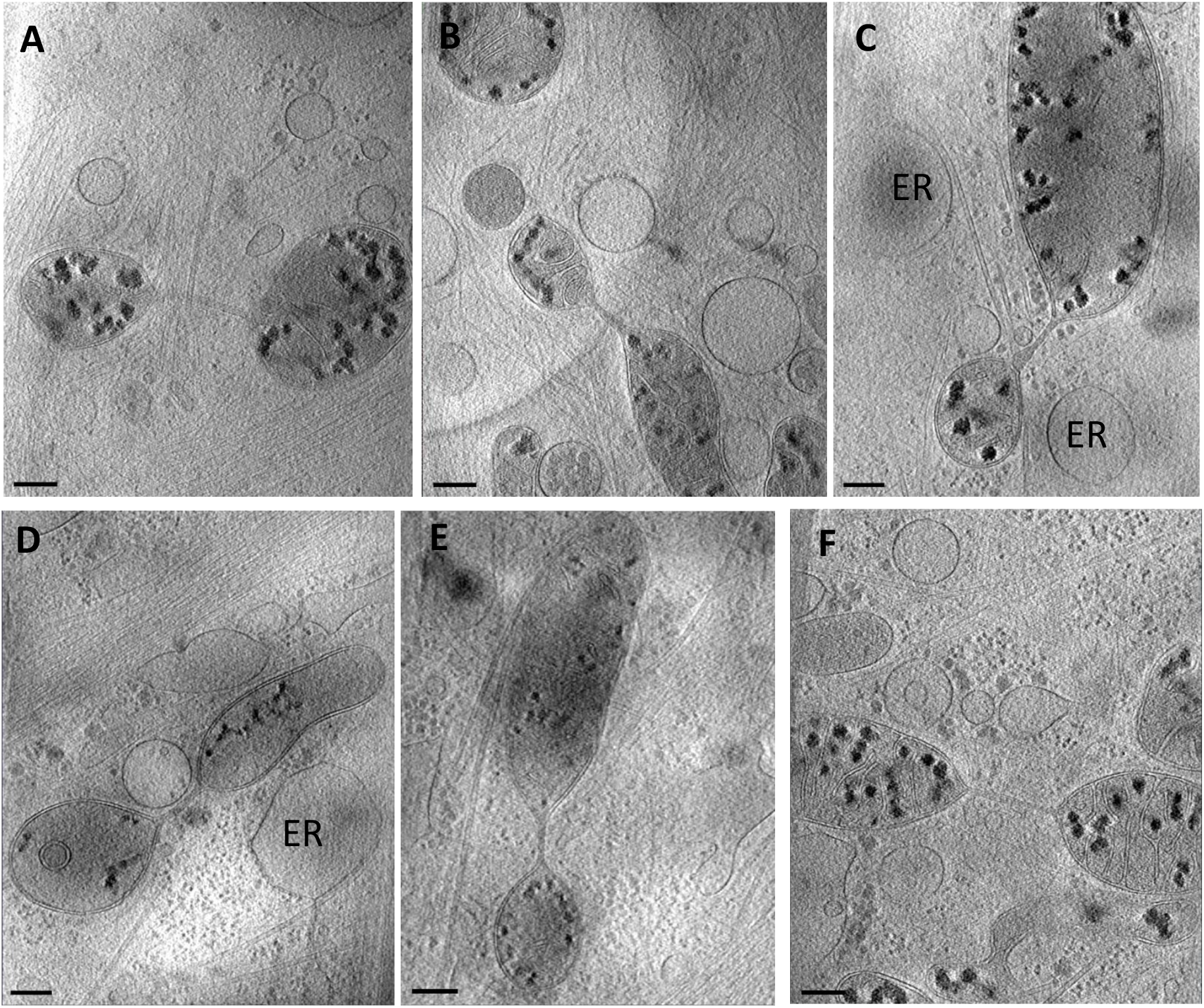
Single mitochondrial budding. 2D slices of 3D tomography reconstructions show individual budding events characterized by a connecting structure between a larger mitochondrion and a smaller mitochondrion. The connecting structures are usually thin and long. One sees no ring, or ring-like structures, or ER wrapping of the bridging region, suggesting that the budding is independent of Drp1 and ER. Bars, 200nm.

**Fig. 5.**
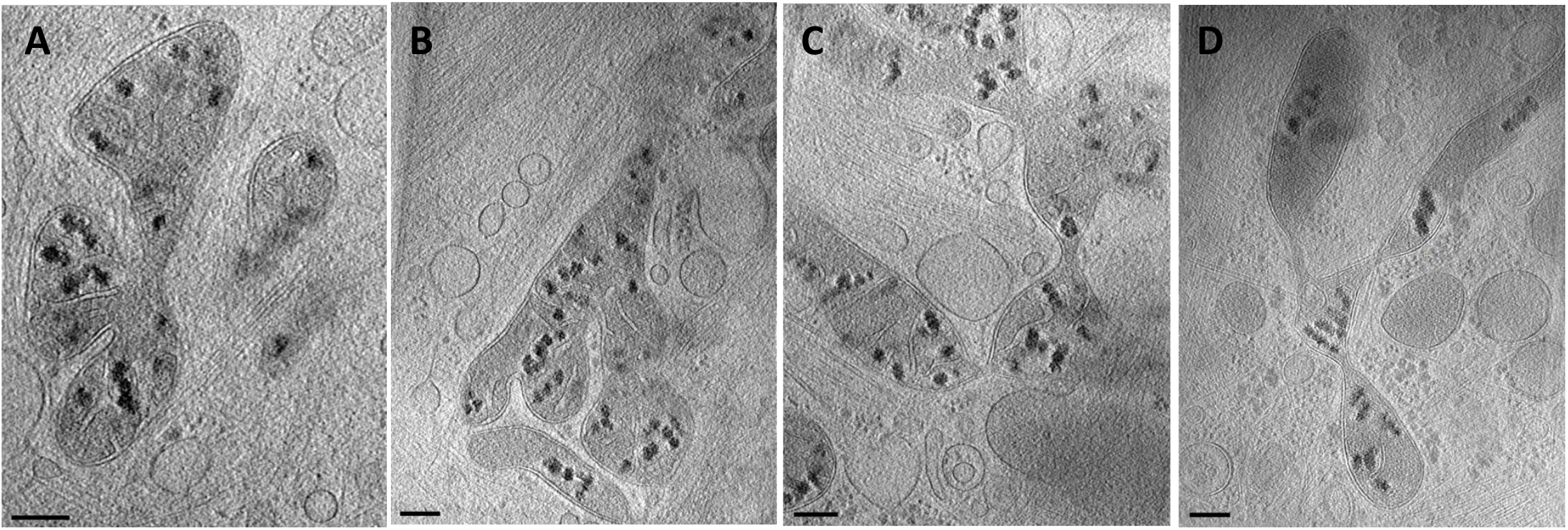
Multiple mitochondrial budding. 2D slices of 3D tomography reconstructions show multiple characteristic budding features. For the first time, we observed flower-like mitochondria, which may represent simultaneous budding or sequential budding from the same mother mitochondrion. Bars, 200nm.

### Structures of multiple mitochondrial budding as shown in Fig.5 have never been reported before

They may represent simultaneous budding of multiple daughter mitochondria or sequential budding of multiple mitochondria from the same mother mitochondrion as previously observed (Hu 2014). Clearly, most of the mitochondrial budding sites observed by *in-situ* Cryo-ET are usually quite long and do not present ER wrapping (Hu 2014, Nedozralova et al. 2022) or ring-like structures that constrict the mitochondrion, suggesting neither ER nor DRP-1 is required for mitochondrial budding. It may also suggest Drp1- and ER-mediated mitochondrial fragmentation is a different process from mitochondrial budding.

The narrow connecting mitochondrial structures may extend to several hundred nanometers Fig.6, as previously observed by super-resolution microscopy visualization (Shim 2012) and Cryo-ET (Hu 2014). The diameter of narrow regions is usually about 50 nm, consistent with previous Cryo-ET results (Hu 2014), but distinct from the 100 nm average diameter observed by fluorescence microscopy (Shim 2012). The discrepancy may be due to the accuracy of the two imaging techniques.

**Fig. 6.**
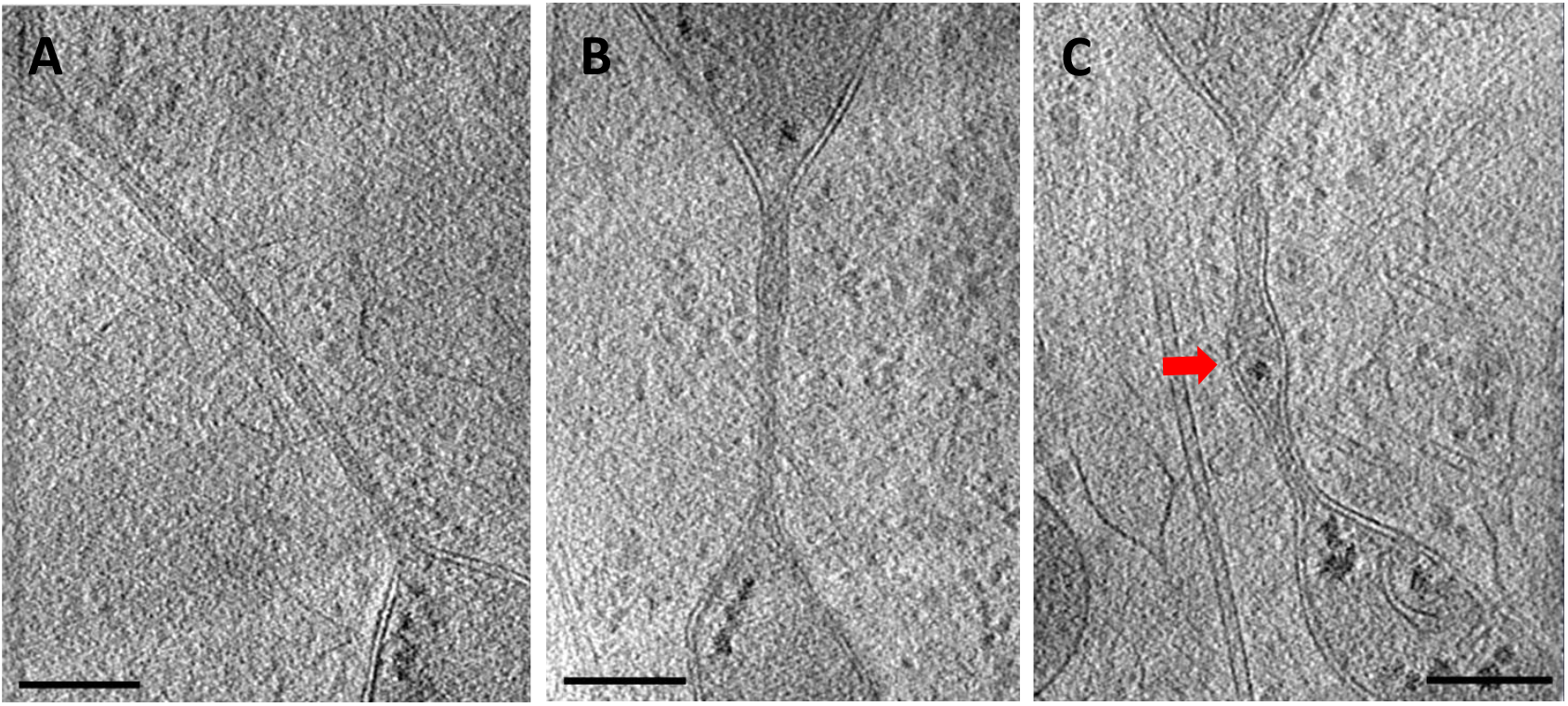
Long bridging mitochondrial structures. Several hundred-nanometer-long connecting structures were observed as in HeLa cells. The typical diameter is about 50 nm but may vary significantly and may even develop into a new mitochondrion (Fig. 6C), contrasting with the report from previous super-resolution microscopy but consistent with a previous Cryo-ET study. Bars, 200nm.

Mitochondrial derived small vesicles were reported in previous studies (Sugiura et al. 2014). Extracellular mitovesicles were also observed in the study of Down syndrome (D’Acunzo et al., 2021). Intra- and extracellular vesicles were abundant in *in-situ* Cryo-ET images as shown in Fig.7A pointed by white arrows. Interestingly, we observed direct interaction/contacts of mitochondria and vesicles as shown in Fig.7B-E. These vesicles differ significantly from previously reported small mitochondrial derived vesicle by size, e.g., over 500 nm in this study vs. 70-100 nm in a previous study (Sugiura et al., 2014). They also differ from the extracellular mitovesicles (D’Acunzo at el., 2021) because they are intracellular. The meaning of these large vesicles that directly interact with mitochondria requires further investigation. Our preliminary hypothesis is that they may have mitochondrial origins.

**Fig. 7.**
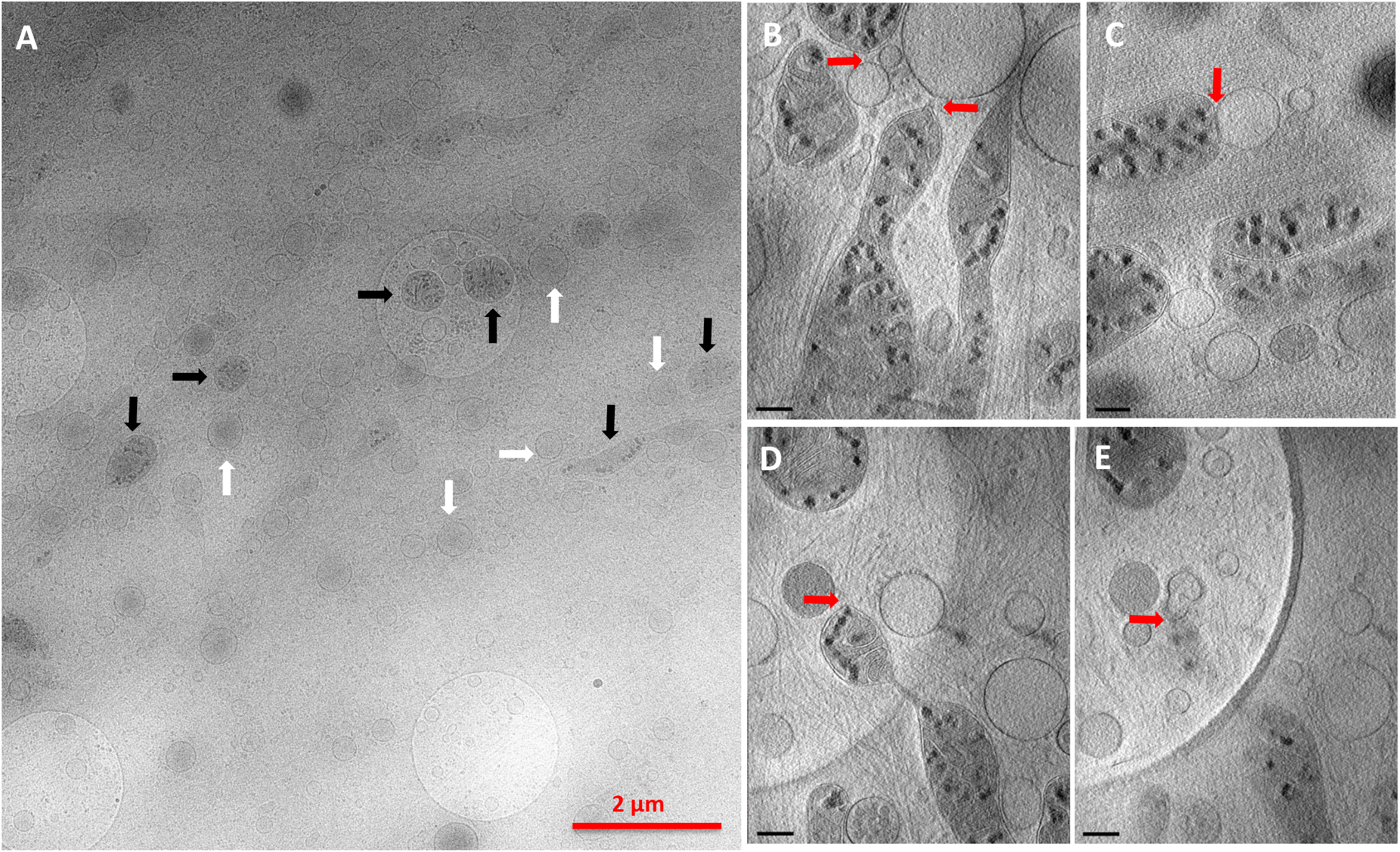
Direct interaction of mitochondria and vesicles. A, Low magnification image showing a cellular region with abundant mitochondria (black arrows) and vesicles (white arrows). Connections (B, D/E) and membrane fusion (C) between mitochondria and vesicles can be appreciated. (D) and (E) are two 2D slices of the same 3D reconstruction at different heights. Together they show a vesicle interacting/associating with a mitochondrion in the vertical direction (Supplement movie). Note that the mitochondrion that is connected to the vesicle is a mitochondrial bud connected to a mother mitochondrion as shown in D. Bars, 200nm, or otherwise stated.

## Conclusion

The large scale of mitochondrial budding structures observed in this study, including single and multiple mitochondrial budding, confirms our previous study in HeLa cells, and provides new evidence to support that mitochondrion budding is the general mechanism for mitochondrial division. This study shows clearly that mitochondrial budding is independent of Drp1 or ER. It indicates the Drp1- and ER-mediated mitochondrial dynamics is a processing different from mitochondrial budding. This study highlights the need of further investigation to identify the key molecular factor(s) that regulates mitochondrial budding.

Meanwhile, we observed direct interactions of mitochondria and large vesicles for the first time. The large vesicles differ significantly from previous reported small mitochondrial-derived vesicles in size and from extracellular mitovesicles by their locations. We hypothesize that those large vesicles may have mitochondrial origins.

## Materials and Methods

### Specimen Preparation

BS-C-1 cells were acquired from ATCC (www.atcc.org) and cultured according to the ATCC protocol, i.e., in growth media of Eagle’s Minimum Essential Medium (EMEM, ATCC) supplemented with 10% Fetal Bovine Serum (FBS, ATCC), and at 37 °C with 5% CO_2_. We glow-discharge-treated 200 mesh R1/4 or R2/4 Quantifoil holey carbon gold grids (Quantifoil^®^, Grossloebichau/Jena Germany) in a Pelco easiGlow (Ted Pella, USA) with 25 mA current for 20 seconds, before sterilization by UV light for 15 minutes on each side in a cell-culture hood. Sterilized grids were placed in a cell-culture petri dish (Corning, Corning, NY, USA) containing complete cell growth media, and incubated in the cell culture incubator at 37 °C with 5% CO_2_ for at least 15 minutes. And then we introduced BS-C-1 cells into the cell culture petri dish. EM grids on which cells grew to the confluence of 50% or above but below 100% were plunge-frozen with a Lecia EM GP2 plunge freezer. EM grids were blotted for 8 seconds on the back side to minimize the disturbance of the cells when freezing. Frozen grids were stored in the liquid nitrogen storage tank until Cryo-EM examination.

### Data Acquisition

We imaged Frozen EM grids on a ThermoScientific Titan Krios G3i Cryo-TEM operated at 300kV and at liquid nitrogen temperature. Tomography images were filtered with a BioQuantum energy filter and acquired with a Gatan K3 Direct Electron Detector (Gatan, Pleasanton, CA, USA). We used Thermo Scientific Tomography program at the nominal magnification of either 33,000x with tilt range of -/+ 48 degree and 3-degree increment, or 19,500x with tilt range of +/-50 degree and 2-degree increment. The dose was a constant 3 e^-^/Å^2^ at each tilt angle for the +/-48 deg. tilt range, and 2.5 e^-^/Å^2^ for the +/-50 degree tilt range. The defocus was set to -5 µm for both tilt schemes.

### Image Processing

The motion correction of the frames was done with MotionCor2 (Zheng et al., 2016). The frames were put back to stack files by using scripts from Dr. Jun Liu’s lab of Yale. The tomography tilt series were aligned and reconstructed using the fiducial-mark-free alignment program AreTomo (Zheng et al., 2022). All 3D reconstructions were denoised and filtered by nonlinear anisotropic diffusion in IMOD (Kremer et al., 1996). Visualization was also run in IMOD.

## Acknowledgement

This project was sponsored in part by Brookhaven National Laboratory’s High School Summer Research Program. The work was supported by the DOE Office of Science, Office of Basic Energy Sciences, and especially the Physical Biosciences program of the Chemical Sciences, Geosciences and Biosciences Division under contract no. DE-SC0012704 (to C-JL). Specimen preparation, including cell culture and cell freezing, was performed using the Bioimaging facility at the NSLS-II of BNL. Data were collected at the Laboratory for Biomolecular Structures of BNL, which is supported by the DOE Office of Biological and Environmental Research (KP1607011).

A manuscript of this work was previously deposited as a preprint on BioRxiv (doi: 10.1101/2023.11.10.566563)

## Author contribution

GH and C-JL conceived the research. GH designed the experiments. JH, XZ, JQ, and GH did cell culture. JH and GH prepared Cryo-EM grids, collected, and processed data, and drafted the manuscript. All the co-authors edited the manuscript.

## Notes

### Competing Interest Statement

The authors have declared no competing interest.

### Summary of Updates

References extended, writing polished.

